# Topological Insights into Mutant Huntingtin Exon 1 and PolyQ Aggregates by Cryo-Electron Tomography

**DOI:** 10.1101/2020.12.08.417048

**Authors:** Jesús G. Galaz-Montoya, Sarah H. Shahmoradian, Koning Shen, Judith Frydman, Wah Chiu

**Affiliations:** Department of Bioengineering and James H. Clark Center, Stanford University, Stanford, CA 94305, United States of America; Department of Biology and Chemistry, Laboratory of Biomolecular Research, Paul Scherrer Institute, Villigen, Switzerland; Department of Biology, Stanford University, Stanford, CA 94305, United States of America; Division of CryoEM and Bioimaging, SSRL, SLAC National Accelerator Laboratory, Menlo Park, CA 94025, United States of America

## Abstract

Huntington disease (HD) is a neurodegenerative trinucleotide repeat disorder caused by an expanded poly-glutamine (polyQ) tract in the mutant huntingtin (mHTT) protein. The formation and topology of filamentous mHTT inclusions in the brain (hallmarks of HD implicated in neurotoxicity) remain elusive. Using cryo-electron tomography and subtomogram averaging, here we show that mHTT exon 1 and polyQ-only aggregates *in vitro* are structurally heterogenous and filamentous, similar to prior observations with other methods. Yet, we observed some filaments in both types of aggregates under ∼2 nm in width, thinner than previously reported, while other regions form large sheets. In addition, our data show a prevalent subpopulation of filaments exhibiting a lumpy, slab-shaped morphology in both aggregates, supportive of the “polyQ core” model. This provides a basis for future cryoET studies of various aggregated mHTT and polyQ constructs to improve their structure-based modeling and their identification in cells without fusion tags.

## INTRODUCTION

Huntington disease (HD) is a neurodegenerative, fatal trinucleotide repeat disorder caused by a polyQ expansion in exon 1 of mutant huntingtin (mHTT) exceeding a pathogenic threshold of Q > ∼35 (MacDonald et al., 1993). Patients suffer from motor and cognitive impairments and despite our increased understanding of HD (Testa & Jankovic, 2019) and promising clinical trials (Tabrizi et al., 2019), cures and preventive treatments remain elusive (Wild & Tabrizi, 2017).

Expression of mHTT exon 1 (a caspase cleavage product within cells, hereafter “mEx1”) elicits HD phenotypes in cellular and animal models (Mangiarini et al., 1996; von Hörsten et al., 2003; Wang et al., 2006; Wang & Sigworth, 2006), including primates (Yang et al., 2008). Furthermore, mEx1 inclusions in mouse and human brains (Davies et al., 1997; DiFiglia et al., 1997) are morphologically similar to those in R6/2 and mEx1 knock-in mice (Sathasivam et al., 2010).

A polyQ expansion in different genes causes at least eight other disorders with a similar pathogenic Q-repeat length threshold, irrespective of flanking motifs (Fan et al., 2014; Gatchel & Zoghbi, 2005), and polyQ peptides as short as Q=20 are toxic when they contain a nuclear localization signal (Yang et al., 2002). Since structure often determines function (Redfern et al., 2008), as shown for mHTT toxic aggregates (Nekooki-Machida et al., 2009; Sun et al., 2015), an increased structural understanding of polyQ aggregates can help uncover the mechanisms underlying their biogenesis, development, and cytotoxicity to better model polyQ disorders.

Both small mHTT oligomers and large inclusion bodies (IBs) can be neurotoxic (Albin, 2017; Nucifora et al., 2001). Filamentous aggregates of mEx1 constructs with various polyQ lengths (mEx1-Qn) have been amply visualized with negative staining transmission electron microscopy (NS-TEM) (Bugg et al., 2012; Crick et al., 2013; Scherzinger et al., 1999), a technique often limited to 2D projections and subject to metal stain and drying artifacts. Two recent studies used cryo focused ion beam milling and electron tomography (cryoFIB-ET) to visualize transfected mEx1-Q97 forming IBs within yeast (Gruber et al., 2018) and HeLa (Bäuerlein et al., 2017) cells. However, a green fluorescence protein (GFP) fusion tag was used, which can alter mEx1 aggregation (Warner et al., 2017).

Here, we used direct observation (without heavy metal stain or fusion tags) by cryo-electron tomography (cryoET) and subtomogram averaging (STA) (Galaz-Montoya & Ludtke, 2017; Zhang, 2019) to visualize vitrified filamentous mEx1-Q51 and Q51 (a peptide consisting of only glutamines) aggregates *in vitro*. We leveraged our initial observations of mEx1-Q51 filaments by cryoET (Darrow et al., 2015; Shahmoradian et al., 2013; Shen et al., 2016) and capitalized on recent algorithmic developments including compressed sensing for tomographic reconstruction (Chen et al., 2017b; Deng et al., 2016), convolutional neural networks for feature annotation (Chen et al., 2017a), and automated fiducial-less tiltseries alignment and subtiltseries refinement for subtomogram averaging (Chen et al., 2019) to resolve previously unattainable structures.

Our study provides a three-dimensional (3D), nanometer-resolution structural description of vitrified mEx1 and Q-only aggregates at an unprecedented level of detail, while avoiding the use of chemical fixatives and confounding fusion tags.

## RESULTS

### Mutant huntingtin exon 1-Q51 filaments exhibit a large variation in width, narrow branching angles, and “lamination”

We analyzed tomographic tiltseries of vitrified mEx1-Q51 **(Figure 1A)**, collected as previously described (see Methods)(Shahmoradian et al., 2013). Owing to the higher contrast and minimized missing wedge artifacts attainable with compressed sensing (CS) compared to standard weighted back projection (Deng et al., 2016), we incorporated CS in our pipeline to reconstruct the tiltseries into tomograms (see Methods), which exhibited aggregated filamentous densities (**Figure 1B**,**C**). While CS might introduce artifacts at very high resolution in the subnanometer range, it has been demonstrated to produce faithful reconstructions at nanometer resolution (Jin et al., 2018). The most frequently observed widths from aggregates in six tomograms ranged between ∼5 and ∼16 nm, with the thinnest filaments exhibiting regions down to ∼2 nm thickness (**Figure 1D**,**E**). On the other hand, the thickest filaments measured over ∼20 nm in width. These measurements are not consistent with a cylindrical shape of a single radius, as reported in recent cryoFIB-ET studies (Bäuerlein et al., 2017; Gruber et al., 2018). Rather, our observations are consistent with a heterogeneous plethora of thin filaments, rectangular prisms, and even “sheets” of varying size. We interpret the predominant species among our observed “filaments” as 3D rectangular “slabs”, which could exhibit many different center-slice widths in between their widest and narrowest dimensions when sliced computationally at slanted angles. Our computational simulations of filamentous subtomograms using EMAN2 (Galaz-Montoya et al., 2015) support this model (**Supplementary Figure S1**).

**Figure 1.**
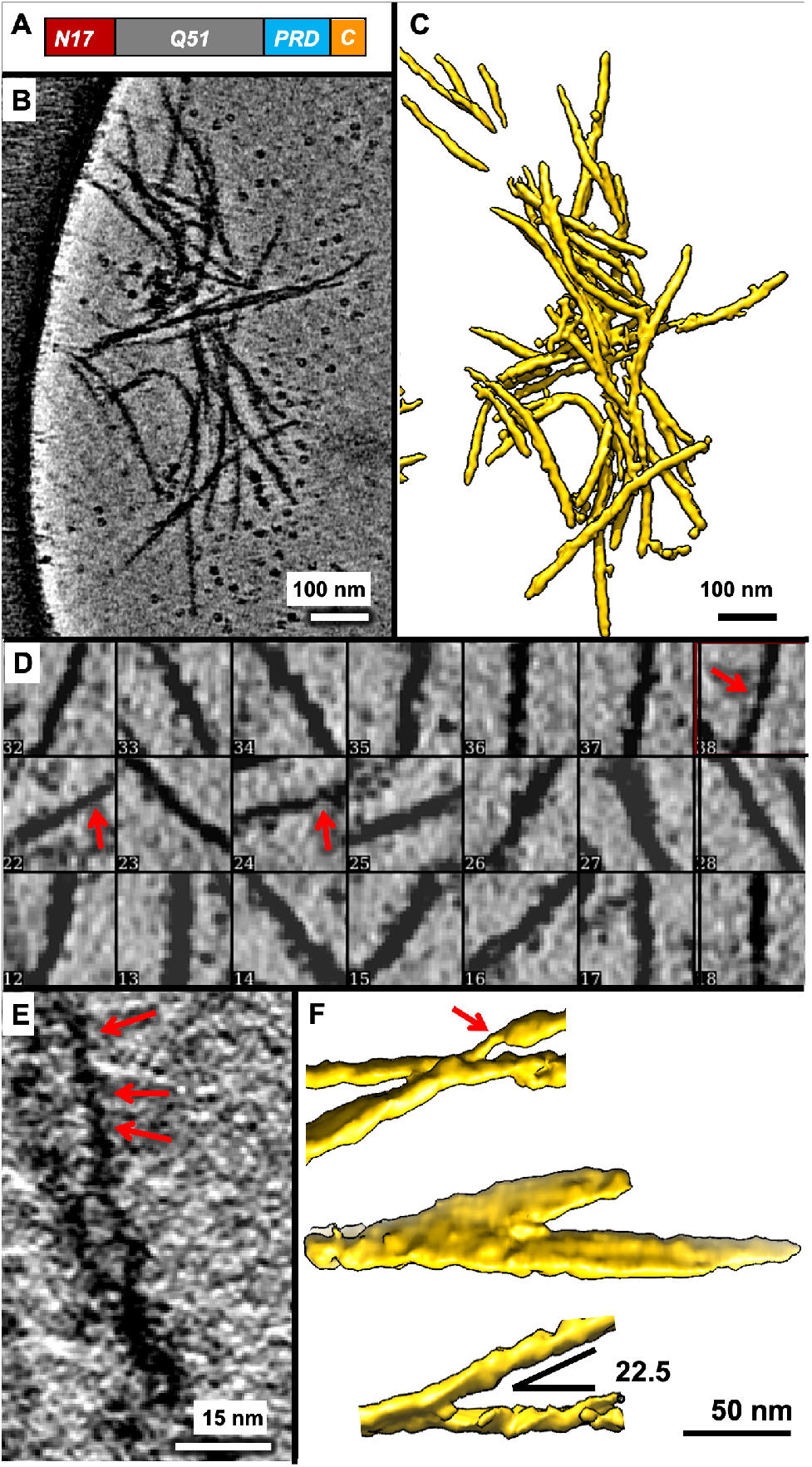
MEx1-Q51 filaments exhibit a very large variation in width within and across filaments. **(A)** Schematic of the mEx1-Q51 construct. **(B)** Slice parallel to xy (∼1.7 nm thick) through a representative 4x down-sampled cryoET tomogram of aggregated mEx1-Q51, reconstructed with compressed sensing, lightly filtered to enhance visualization, and **(C)** corresponding semi-automated 3D annotation. **(D)** Selected areas from slices of large mEx1-Q51 aggregates showing individual filaments, widely varying in width, with the thinnest filaments exhibiting regions down to ∼2 nm width, indicated by the red arrows. **(E)** Zoomed-in view of a xy slice (∼0.4 nm thick) from a selected region of a tomogram without any down-sampling, showcasing ultra-thin regions in mEx1-Q51 filaments. **(F)** Sections of annotated mEX1-Q51 filamentous aggregates from cryoET tomograms showing relatively narrow branching angles and an example of a thicker “laminated” sheet-like region (the annotation example in the middle).

We used semi-automated annotation based on neural networks (Chen et al., 2017a) to visualize in 3D what kind of objects yielded the extensive width variations detectable in 2D slices of 3D tomograms. Visualizing mEx1 filaments as 3D isosurfaces (**Figure 1C**; **Supplementary Figure S2**) revealed filaments of different dimensions altogether, including regions that appeared as “sheets” as thick as ∼50 nm, estimated from the annotations and from their persistence through 2D slices. The mEx1 filaments seemed to predominantly branch out at angles varying from ∼10° to ∼45° (only sporadically larger), with angles between ∼20° to ∼25° being most common (**Figure 1F**).

### Subpopulation of mEx1-Q51 filament segments exhibits a lumpy, slab-shaped morphology

Many filaments appeared to be “lumpy” both in 2D slices from 3D tomograms (**Figure 2A**) as well as in in 3D annotations (**Figure 2B**), suggestive of potential periodicity. Thus, we performed subtomogram averaging (STA) of filament segments, avoiding obviously-laminated regions and thick bundles. The subtomogram average of mEx1-Q51 filament segments (n=450, from 6 tomograms) converged to a lumpy ∼7×15 nm slab at ∼3.5 nm resolution (**Figure 2C**). The Fourier transform of 2D re-projections of the average did not reveal crisp layer lines, in agreement with previous studies suggesting that mEx1 filaments do not exhibit a canonical amyloid structure with parallel subunits stacked helically in register (Bugg et al., 2012). Indeed, HD does not strictly fit among the diseases known as “amyloidoses” (Dobson, 2001); nonetheless, the power spectra showed bright maxima off of the meridian, at ∼11.7 nm (**Supplementary Figure 4A**), suggestive of potential periodicity for at least relatively short stretches (∼65 nm, the length included in the extracted subtomograms).

**Figure 2.**
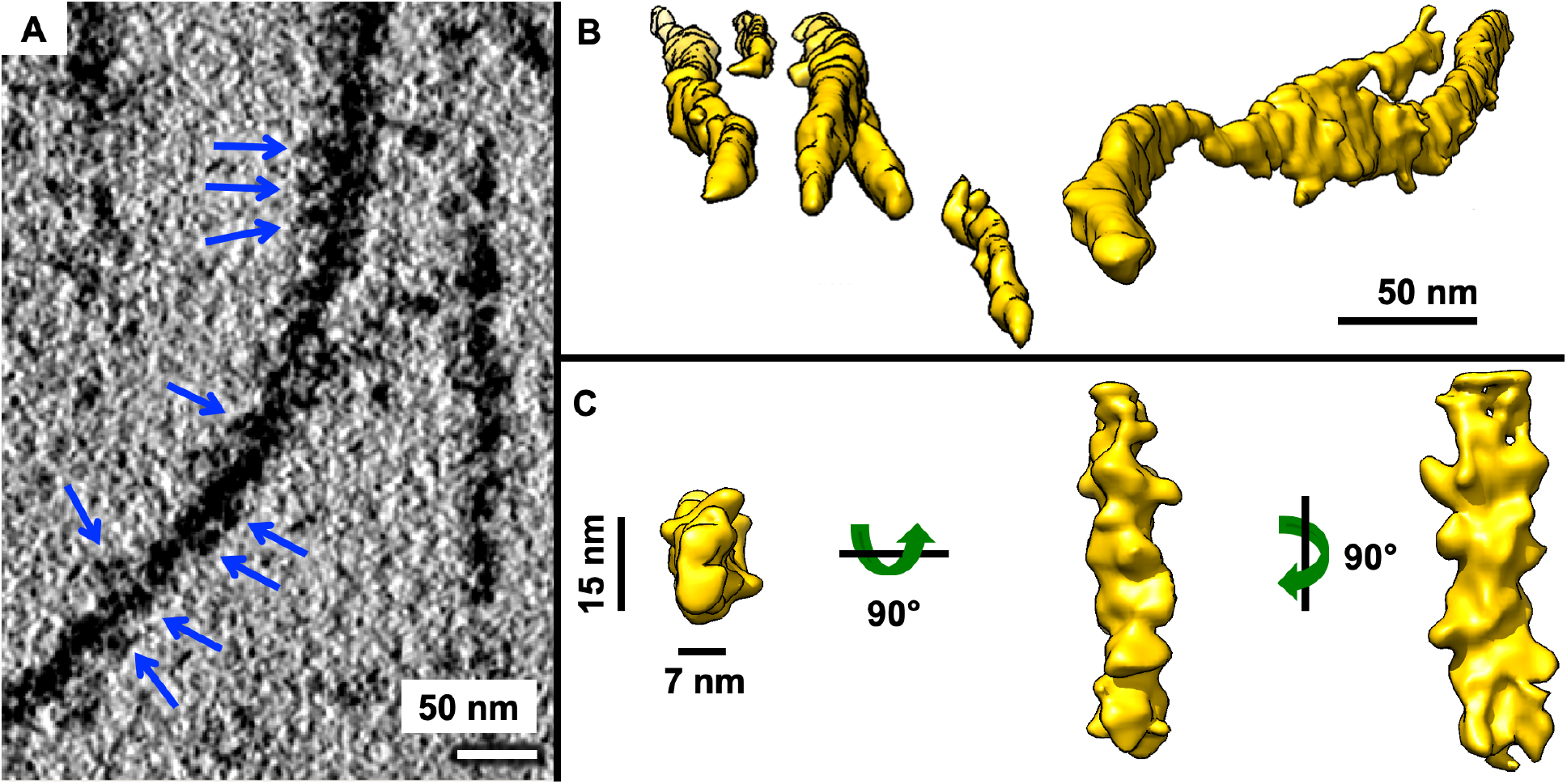
Aggregated mEx1-Q51 exhibits lumpy, slab-shaped filaments. **(A)** Pseudo-periodic pattern of repeating “lumps” (blue arrows) along the length of an mEx1-Q51 filament as seen in an xy slice (4.4 Å thick) from a tomogram of aggregated mEx1-Q51. **(B)** Selected regions from semi-automated neural network annotations showing lumpy filaments of various widths, including sheet-like regions (middle region of right-most example). **(C)** Subtomogram average of a subpopulation of filament segments exhibiting a lumpy 7×15 nm slab-shaped morphology.

### Lumpy, slab-shaped Q51-only filaments also exhibit lamination

Since an expanded polyQ tract is the common culprit of all polyglutamine diseases, we performed the same analyses for a Q51-only peptide (**Figure 3A**) as reported above for mEx1-Q51. We found that Q51 also forms aggregates (**Figure 3B, Supplementary Figure 3**) exhibiting lumpy filaments (**Figure 3C**) of varying widths (**Figure 3D**,**E**), with regions as thin as ∼2 nm.

**Figure 3.**
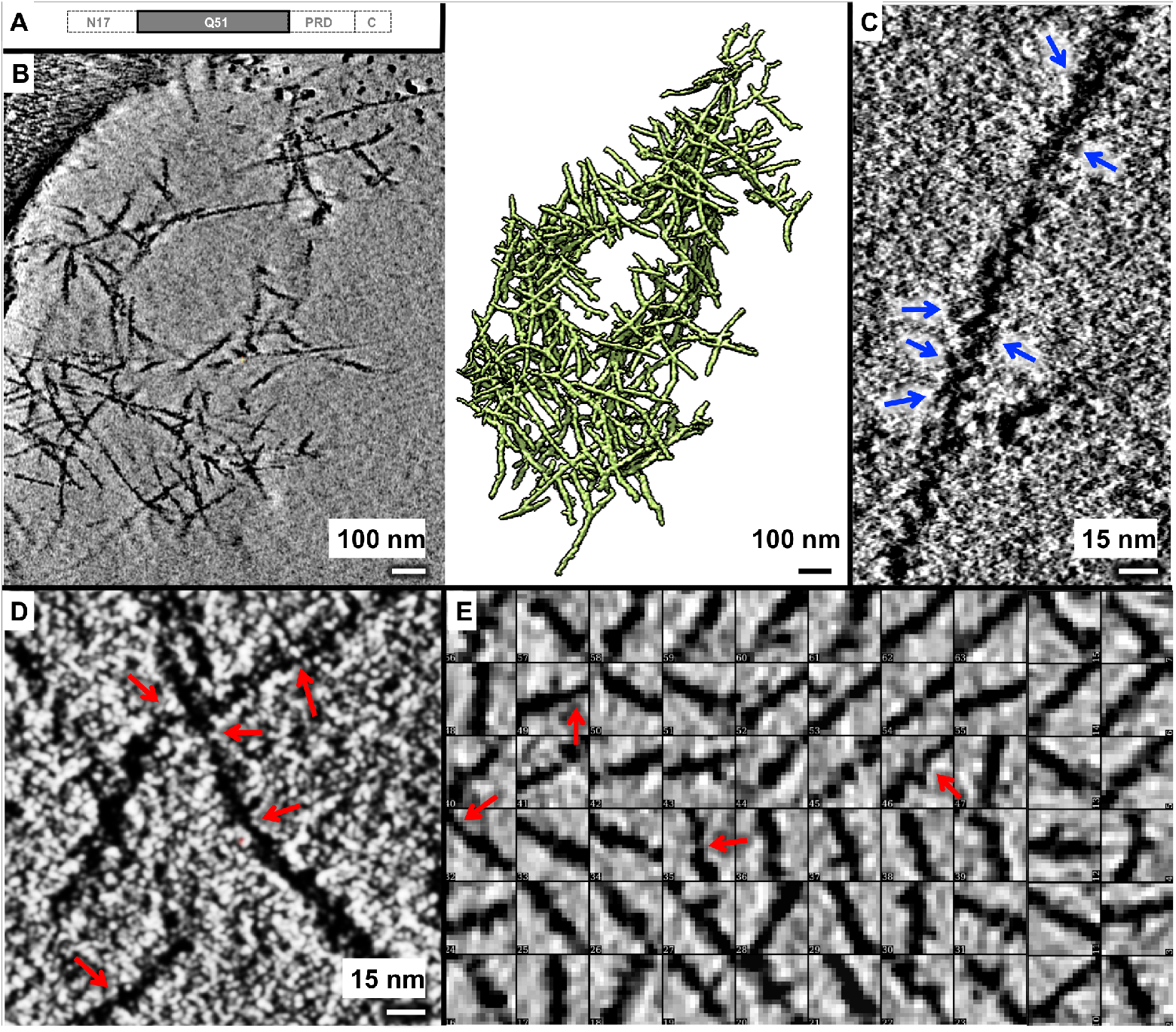
Lumpy Q51 filaments exhibit a large range of widths. **(A)** Schematic of the Q51 construct, lacking all mEx1 domains except for the polyQ tract. **(B)** Slice parallel to the xy plane (∼2.1 nm thick) through a representative 4x down-sampled cryoET tomogram of aggregated Q51 reconstructed with compressed sensing (left) and corresponding 3D annotation (right). Zoomed-in views of xy slices (∼0.5 nm thick) from selected regions of the tomogram shown in A but without any down-sampling, exhibiting **(C)** a pseudo-periodic pattern of repeating “lumps” along the length of a Q51 filament (blue arrows), and **(D)** regions in thin filaments that are as thin as ∼2 nm in width (red arrows). **(E)** Examples of 2D xy slices through representative 3D subtomograms of Q51 filaments showing a wide variation in width, including super-thin regions ∼2 nm in width (red arrows).

Q51 filaments branched out/crossed over more often and at wider angles than mEx1-Q51 filaments, with ∼60° being the most common angle (**Figure 4A**). Furthermore, Q51 filaments exhibited larger lamination sheets than those of mEx1-Q51 of up to ∼60+ nm in width (**Figure 4B**). The subtomogram average of non-laminated filament segments (n=493, from 6 tomograms) converged to ∼3.2 nm resolution and also revealed a lumpy ∼7×15 nm slab (**Figure 4C**), with a crossover length of ∼11.2 nm according to the power spectrum of its projections (**Supplementary Figure 4B**), all strikingly similar results to those obtained for mEx1-Q51.

**Figure 4.**
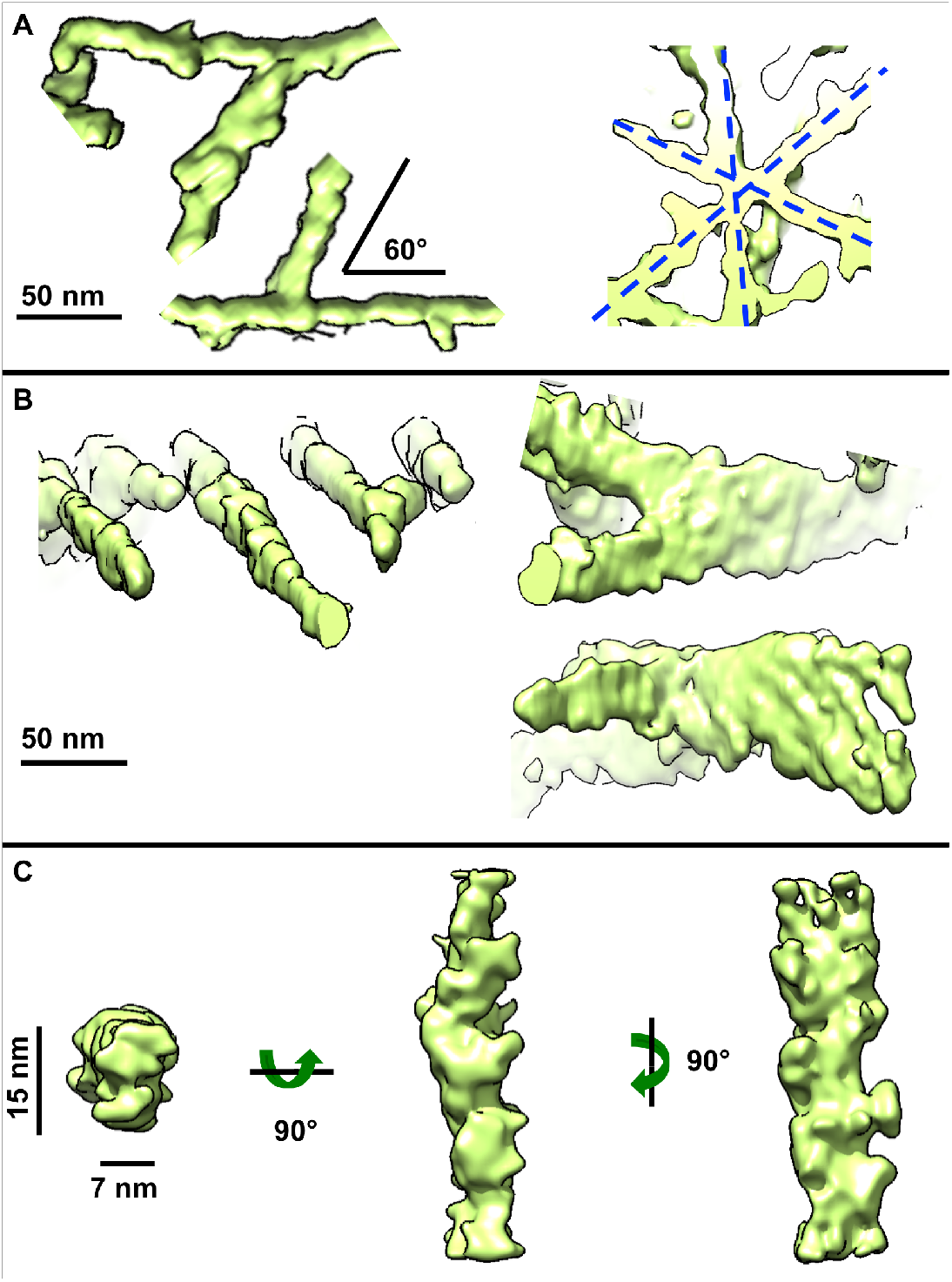
Aggregated Q51 exhibits lamination sheets and predominantly lumpy, slab-shaped filaments. **(A)** Representative sections of annotated Q51 filamentous aggregates from cryoET tomograms showing their most common branching/crossover angle (∼60°), often in an asterisk-like pattern, and **(B)** thicker regions (right) akin to lamination, onside thinner ones (left). **(C)** Subtomogram average of a subpopulation of filament segments exhibiting a lumpy 7×15 nm slab-shaped morphology.

## DISCUSSION

In one of the earliest reports visualizing mEx1-Q51 filamentous aggregates by NS-TEM, filaments digested with factor Xa or trypsin, which removes mEx1’s N17 domain critical to mHTT localization and function (Maiuri et al., 2013), were reported to have a diameter of ∼7.7-12 nm from 2-dimensional (2D) images (Scherzinger et al., 1997). These were occasionally referred to as “ribbons”, and other 2D NS-TEM observations have reported similar filaments with a 10-12 nm “diameter”, which may associate laterally (Bugg et al., 2012). However, apparent lateral associations in 2D NS-TEM observations may arise from the compression of all densities into a single layer due to dehydration. In contrast, the vitrified filaments seen in our 3-dimensional (3D) tomograms varied much more in width within and across filaments, similar to those reported in studies visualizing aggregated Q-only peptides with NS-TEM (Chen et al., 2002; Chen et al., 2001), which also detected wide ribbons and thin filaments under different incubation temperatures and using a “freeze-concentration” method involving cycles of freezing and thawing. Here, our observations did not suffer from NS-TEM artifacts and were conducted at 30° C, without special temperature manipulations.

While other amyloidogenic filaments have been amply studied using cryo electron microscopy (cryoEM) (Fitzpatrick & Saibil, 2019; Fitzpatrick et al., 2017; Gremer et al., 2017; Guerrero-Ferreira et al., 2018; Iadanza et al., 2018; Li et al., 2018a; Li et al., 2018b; Scheres et al., 2020), cryoEM studies of mHTT and polyQ-containing aggregates have been extremely scant in comparison, likely due to the extensive conformational heterogeneity of these specimens (Wetzel, 2012), which limits the applicability of “bulk techniques” (*e*.*g*., Circular Dichroism) and calls for the increasing application of “single molecule” techniques (Ruggeri et al., 2016). Single molecule techniques such as Atomic Force Microscopy (AFM) and various modalities of electron microscopy (EM) can observe individual components in aggregates (molecules, oligomers, filaments). For EM-related methods, these components can be classified prior to averaging. Of note, cryoET should be the method of choice for relatively thick samples exhibiting extensive conformational and compositional heterogeneity as it avoids adsorption of the specimen onto 2D surfaces and the confounding effect of potentially overlapping densities from different components, as in 2D projections produced by single particle cryoEM.

In two recent cryoFIB-ET studies, mEx1-Q97-GFP filaments were observed and annotated in transfected cellular systems but were not averaged. Rather, they were either modeled as cylinders with an 8-nm diameter for template-based annotation (Bäuerlein et al., 2017) or were segmented as 16 nm filaments (Gruber et al., 2018), surprisingly twice as thick in the latter study than in the former, perhaps owing to differences in the non-native expression systems used or to the confounding presence of GFP fusion tags. Indeed, there can be caveats to using fusions to fluorescent proteins as tags, from impairing the viability and growth of cells via toxic effects from tag aggregation, excitation, or photoactivation, to changing the structure, function, and cellular localization of the tagged protein (Jensen, 2012). Template-based approaches have been successfully applied to annotate more regularly-shaped biological filaments (Rigort et al., 2012; Rusu et al., 2012); however, our data here suggest that the use of a cylindrical template is not an adequate approach to annotate widely heterogeneous mEx1 and polyQ aggregates with filamentous densities of varying dimensions. Indeed, when identifying features in tomograms, template matching can be biased (Yu & Frangakis, 2014) and manual human annotation is subjective and therefore often uncertain and inconsistent (Hecksel et al., 2016). On the other hand, here we used template-free, semi-automated annotation based on machine learning since it can ameliorate these issues by minimizing human input and the use of *a priori* constraints inherent in templates (Chen et al., 2017a).

A recent atomic force microscopy (AFM) study on mEx1-Q49 aggregation suggested that nucleated branching from filaments, rather than lateral associations among them, leads to large bundles (Wagner et al., 2018). However, branching does not explain how the thinner (< 2 nm thick) filaments that we observe here would assemble into thicker slabs and sheets without associating laterally or growing transversally to the main filament axis. Rather, our results suggest that preformed thin filaments can associate laterally and/or that growing filaments can expand transversally in addition to longitudinally, akin to the “lamination” observed for Aβ (Lu et al., 2003), for both mEx1-Q51 and Q51.

While AFM is limited to ∼30 nm in lateral resolution of surface measurements of specimens that are often absorbed and dried onto a 2D substrate, an earlier AFM study of aggregated Q44 peptide detected regions in filament tips with a “height” as thin as ∼5 nm (Poirier et al., 2002). This and the thinness of some of the filament regions we observed here (as thin as ∼2 nm) seem to disagree with the minimum width of ∼7-8 nm proposed for polyQ filaments from various Qn constructs in a prior NMR study that also presented NS-TEM images (Schneider et al., 2011). However, the latter study reports filament widths for a Q54 peptide from NS-STEM images from ∼7-8 nm up to ∼16 nm, in striking agreement with the “short” and “long” sides of the slab-shaped model we propose here as the predominant morphology for filaments formed by both mEx1-Q51 and Q51.

The morphological characteristics deviating from thin cylindrical shapes to form lumpy slabs and sheets may serve as a structural hallmark to identify untagged mHTT aggregates in cells. Furthermore, the more frequent and wider-angle branching of Q-only filaments compared to mEx1 is consistent with our prior 2D cryoEM observations (Shen et al., 2016), suggesting that the N17 domain promotes inter-filament bundling. Conversely, the occurrence of branching might be primarily polyQ-driven. Indeed, our 3D observations here, which are free from fusion tags, stains, dehydration, flattening, and crystallization artifacts, provide a transforming complement and clarification to previous studies by NS-TEM and AFM, as well as light microscopy (Duim et al., 2011; Duim et al., 2014), which visualized mEx1 filamentous aggregates at a coarser level: features often described as “globules” or “oligomers” or “thick filaments” actually correspond to bundles of many interwoven thinner filamentous densities when viewed by cryoET.

The fact that the predominant populations for both mEx1 and Q-only filaments exhibit a similar lumpy slab shape and distance between putative crossovers as revealed by subtomogram averaging suggests that the morphology of their “core” is dictated by and primarily comprised of the polyQ tract, and that the flanking domains in mEx1 are largely exposed at the filament surface, allowing them to modulate inter-filament aggregation. This interpretation agrees with previous nuclear magnetic resonance (NMR) studies on non-pathogenic (Sivanandam et al., 2011) and pathogenic (Hoop et al., 2014; Hoop et al., 2016) mEx1 variants that propose the existence a dense polyQ core.

In one of the latest studies supporting the polyQ-core model (Lin et al., 2017), the authors observed mEx1-Q44 filaments formed at two different temperatures by 2D NS-TEM images (presented in the supplement). The widths reported for these filaments were ∼6.5 nm and ∼15.2 nm, in striking agreement with the dimensions of our slab-shaped subtomogram averages of filament segments from 3D cryoET tomograms of vitrified mEX-Q51 and Q51. While their hypothesis that the thicker ∼15.2 nm filaments must arise from two interwinding protofilaments ∼6.5 nm thick seems to be compatible with our observations here, their model proposing that the flanking domains mediate such interwinding does not explain our observation that polyQ-only filaments also yield a dominant subpopulation with the same ∼7×15 nm slab morphology, which could also correspond to two interwoven protofilaments without flanking domains to bind them. If, indeed, the mEx1-Q51 and Q51-only predominant subpopulations of ∼7×15 nm filaments are composed of two thinner interwinding protofilaments, our data suggest that they might be bound primarily via polyQ-polyQ interactions.

Our observations here warrant further cryoET experiments with much larger datasets of aggregation-competent mEx1 and polyQ-only constructs devoid of even solubilization and purification tags, as even these can cause modest alterations in aggregation kinetics (Adegbuyiro et al., 2017; Vieweg et al., 2016). Datasets at higher magnification and contrast, using state-of-the-art instrumentation, could test whether there exist filament species even thinner than the ∼2 nm regions we observed here, as well as the effects of increasing polyQ length on the 3D morphology of vitrified filaments. Probing the effects of post-translational modifications (PTMs) on filament and overall aggregate morphology with cryoET might be of particular significance, as some PTMs have been found to be modulate aggregation in vivo, with neuroprotective effects (Hegde et al., 2020). Finally, sonication concomitant with trypsin digestion of mEx1 filaments might yield a homogenous-enough population of the polyQ core that may be more amenable to higher-resolution cryoEM/ET studies.

## METHODS

### *In vitro* mEx1-Q51 and Q51 peptide aggregation assays and cryoET sample preparation

We used mutant huntingtin (mHTT) exon 1 with 51 glutamine repeats (mEx1-Q51) and a polyQ-only peptide with 51 repeats (Q51), each of them fused to a TEV cleavage sequence and a GST tag, as previously described (Shen et al., 2016). Aggregation was initiated separately at a concentration of 6 μM for each construct *in vitro* by addition of AcTEV™ protease (Invitrogen), as previously described for mEx1-Q51 (Shahmoradian et al., 2013). The samples were incubated at 30 °C before vitrification. Aliquots of 2.5 μm were separately applied to 200-mesh holey carbon Quantifoil copper grids (previously washed with acetone, and rinsed in PBS) between 4 and 6 h post-initiation of aggregation. The grids were plunge-frozen in a liquid ethane bath kept at liquid nitrogen temperature using a Vitrobot Mark III (FEI Instruments).

### Tiltseries collection

We collected six tiltseries of the Q51 peptide using SerialEM software (Mastronarde, 2005) on a JEM2100 electron microscope operated at 200 kV from -60° to 60° in 2° increments, at 6 μm target underfocus, 5.29 Å/pixel sampling size, with a cumulative dose of ∼80 e/Å^2^. We also reanalyzed a previous dataset comprised of 20 tiltseries of mEx1-Q51 + TRiC, collected similarly to the Q51 peptide dataset, as previously described (Shahmoradian et al., 2013), with a slightly finer sampling size of 4.4 Å/pixel.

### Tomographic reconstruction

All mEx1-Q51 and Q51 tilt series were binned by 2 and initially aligned and reconstructed into tomograms with IMOD (Kremer et al., 1996). Images with artifacts (grid bars in the field of view blocking large regions of the specimen, evident large drift, obvious radiation damage, etc.) were manually removed prior to tiltseries alignment and tomographic reconstruction with weighted back projection. After assessing sample thickness, the tiltseries were reconstructed again into tomograms using compressed sensing (CS) as implemented in ICON-GPU (Chen et al., 2017b; Deng et al., 2016) to improve contrast. Of note, CS also partially restores information that is lacking due to the “missing wedge” artifact inherent in all conventional single-axis limited-angle tomography experiments, such as conventional cryoET (Radermacher, 1988). The tiltseries were aligned and reconstructed yet a third time for subtomogram averaging purposes (as described below), using a new pipeline for cryoET in EMAN2 (Chen et al., 2019) that performs “sub-tiltseries refinement”, akin to prior “hybrid” methods combining concepts from single particle analysis cryoEM and subtomogram averaging (Bartesaghi et al., 2008; Iwasaki et al., 2005; Zhang & Ren, 2012). We processed the mEx1-Q51 and Q51 datasets separately in virtually identical ways.

### Tomogram annotation

Since the ultimate goal of the new EMAN2 cryoET pipeline (Chen et al., 2019) is to perform subtiltseries refinement for subtomogram averaging, tomogram quality only needs to be sufficient to allow for particle identification. Indeed, in EMAN2 not as many parameters are refined during tomographic reconstruction as compared to IMOD, often resulting in lower-quality tomograms. For this reason, we performed all tomographic annotations on better-quality tomograms aligned with IMOD and reconstructed with CS, as described above. MEx1-Q51 and Q51 annotations were carried out on binned-by-4 tomograms using EMAN2’s neural network semi-automated annotation tools (Chen et al., 2017a), except that ∼2-3x as many “references” as the 10 recommended were segmented to seed annotation, and ∼2-3x as many “negative” samples as the 100 recommended were selected to minimize false positives. We initially performed annotation of all mEx1-Q51 and Q51 tomograms by applying the convolutional neural network from the best tomogram to all the rest, separately for each specimen. However, false positives (such as annotating the carbon-hole edge and/or gold fiducials) were reduced further when we generated a neural network specific for each mEx1-Q51 and Q51 tomogram.

### Fibril width range measurements

In all limited-angle tomography experiments (when you cannot tilt through the entire full range from 0° to 180° or -90° to +90° to collect a full set of projections around the object of interest), the “missing wedge” artifact worsens the resolution of raw tomograms along the Z-axis compared to that in the X and Y directions, often giving the appearance of “elongation” of features along the axis with lowest resolution. Therefore, filament widths cannot be accurately measured in 3D from raw tomograms nor their corresponding annotations in arbitrary orientations. The most conservative measurements in the absence of averaging should be performed on slices along the Z-axis of reconstructed tomograms (*i*.*e*., on sections parallel to the XY plane) since features are much less well-resolved in the XZ and YZ planes. Here, we boxed out filament segments for STA (below) and manually measured the thinnest and thickest parts of segments (N ∼100) from the central XY slice of the corresponding subtomogram. The mEx1-Q51 and Q51 data were separately processed in identical ways.

### Initial model generation for subtomogram averaging (STA)

To carry out sub-tiltseries refinement, the new EMAN2 cryoET pipeline (Chen et al., 2019) requires that all steps (from initial tomographic reconstruction) be performed in EMAN2. However, as explained above, whole-tiltseries alignment with IMOD is often superior in quality, given its refinement of more reconstruction parameters, and reconstruction with ICON-GPU can yield higher-contrast tomograms with minimized missing wedge artifacts. Therefore, to generate an initial model, we manually extracted filament segments without much overlap from the best tomogram for each specimen (n=97 for mEx1-Q51; n=135 for Q51) avoiding branching points and regions of dense bundling or obvious lamination. Then, we aligned these subtomograms to a cylindrical reference with a soft edge and computed the average using the legacy tools for STA in EMAN2 (Galaz-Montoya et al., 2016). This average of vertically-aligned filaments was then refined constraining the angular search in altitude to only allow for slightly-tilted orientations (since all particles were already pre-aligned to a cylinder) and flips of 180° in altitude (the other two Euler angles were completely unconstrained). Alignment converged in ∼4-5 iterations for both datasets. We used these preliminary averages as initial models for subsequent unconstrained gold-standard subtomogram averaging of mEx1-Q51 and Q51 with sub-tiltseries refinement in the new EMAN2 pipeline.

### Subtomogram averaging

Since the reconstruction geometry is different for tomograms produced with different software packages, we had to pick subtomograms of filament segments (with < ∼50% overlap) manually from scratch (n=450 for mEx1-Q51; n=493 for Q51) in EMAN2-reconstructed tomograms. Gold-standard refinements seeded with the initial models described above converged in ∼4-5 iteration and ∼60% of the best-correlating particles were kept in the final average for each dataset. The subtiltseries refinement step alone improved the resolution drastically by ∼10 Å or more for both datasets, yielding averages at ∼3.5 nm and ∼3.2 nm resolution for mEx1-Q51 and Q51, respectively, according to the gold-standard FSC=0.143 criterion.

## ACCESSION NUMBERS AND DATA ACCESSIBILITY

The Electron Microscopy Data Bank accession numbers for the structures reported in this paper are as follows: mEx1-Q51 subtomogram average, EMD-21248; Q51 subtomogram average, EMD-21253. The raw data can be made accessible upon request.

## AUTHOR CONTRIBUTIONS

All authors planned and designed experiments. K.S. generated and purified the mEx1-Q51 and Q51 constructs. S.H.S. collected the cryoET data. J.G.G.M. performed all data processing and analyses, prepared all figures, and wrote the manuscript with feedback from all authors.

## ACKNOWLEDGEMENTS AND FUNDING

This research has been supported by grants from the National Institutes of Health, USA, No. NS092525 to J.F. and W.C., and No. P41GM103832 to W.C.

## SUPPLEMENTARY DATA

**Supplementary Figure S1.**
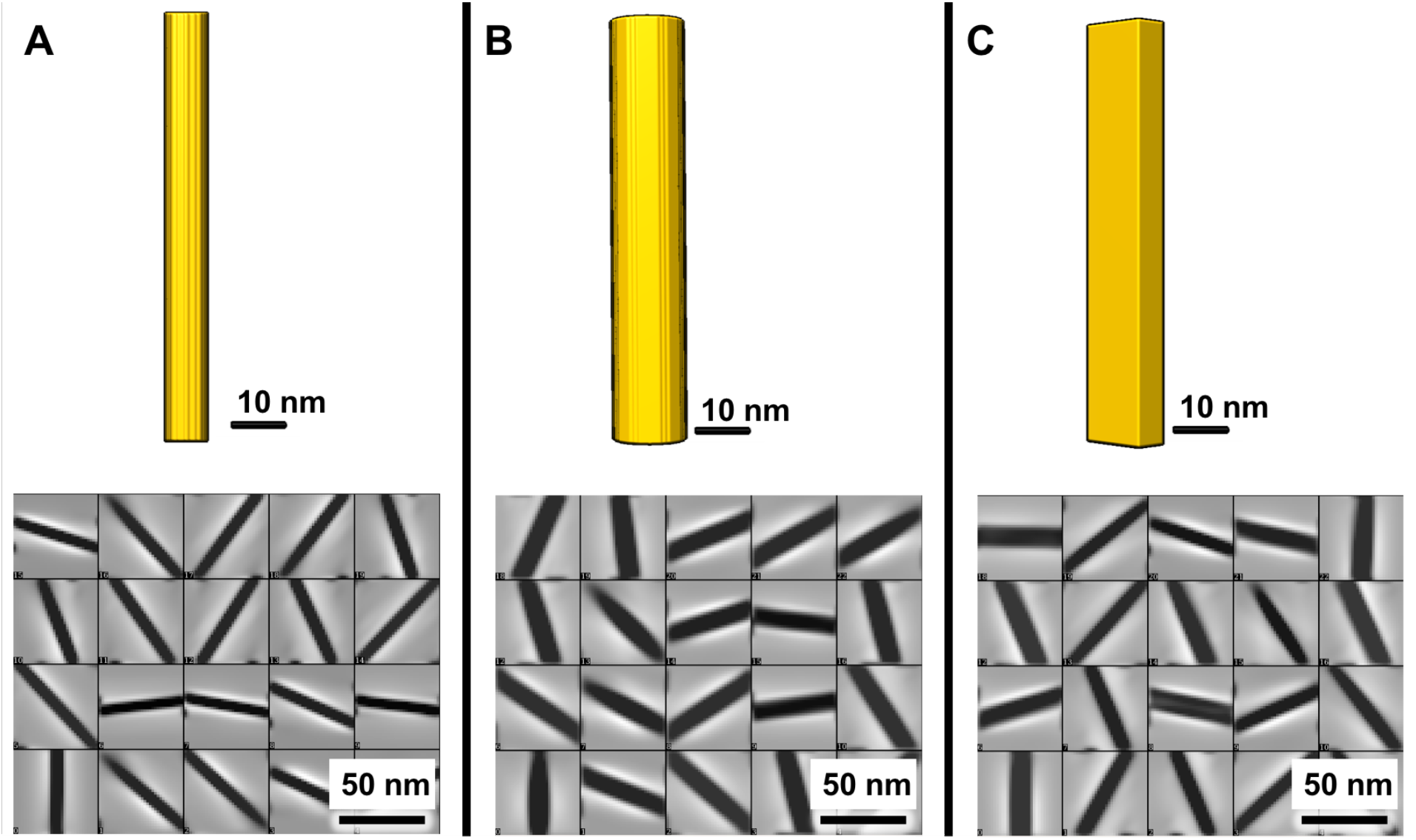
Slab-shaped filaments are more consistent with observations of variable width in central Z cross-sections (in the XY plane, unaffected by the “missing wedge” artifact) than cylindrical filaments. Simulated model and corresponding central Z cross-sections of simulated subtomograms for a cylinder **(A)** 7 nm or **(B)** 15 nm in diameter, and **(C)** a rectangular slab with narrow and wide sides measuring 7 and 15 nm, respectively.

**Supplementary Figure S2.**
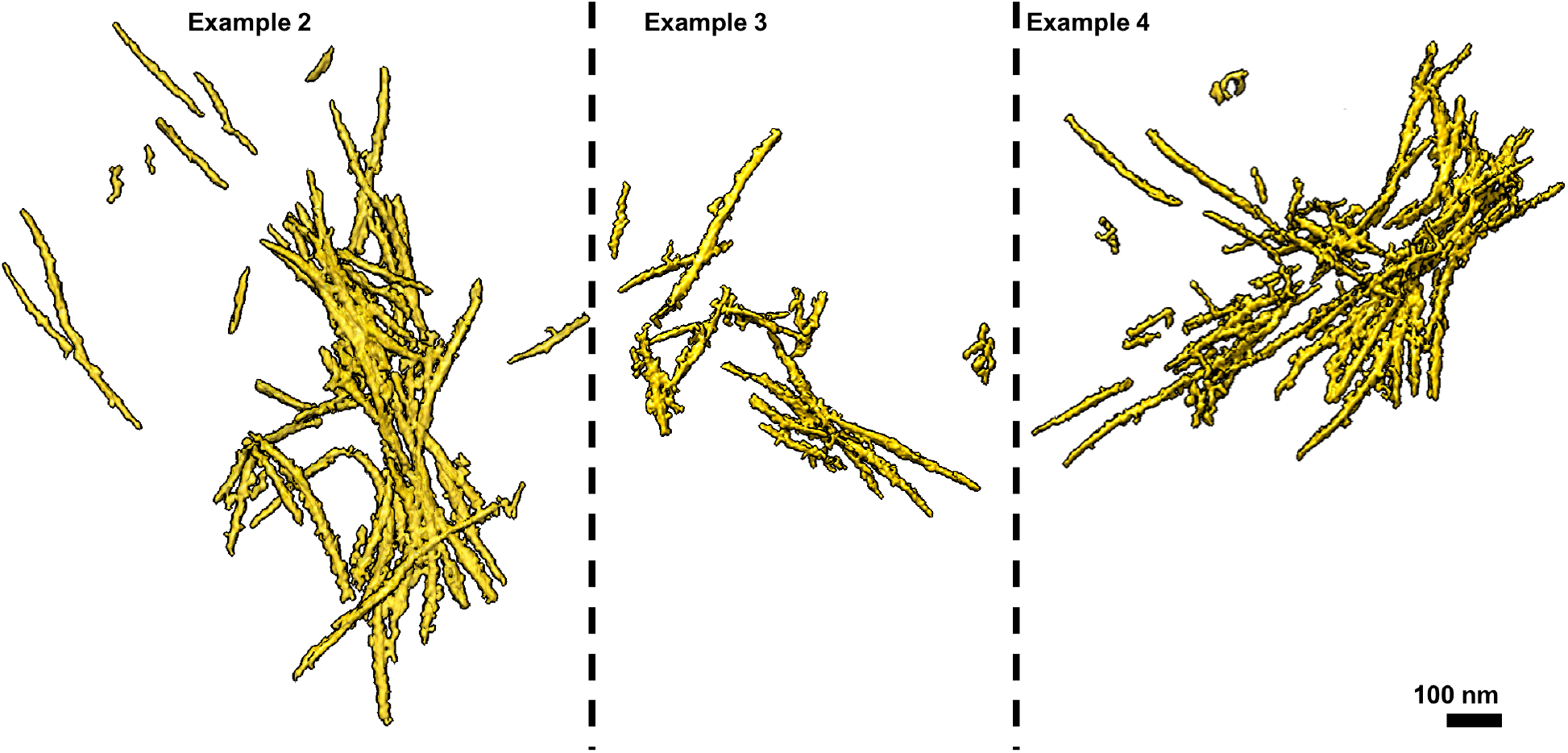
Additional examples of mEx1-Q51 filamentous aggregates.

**Supplementary Figure S3.**
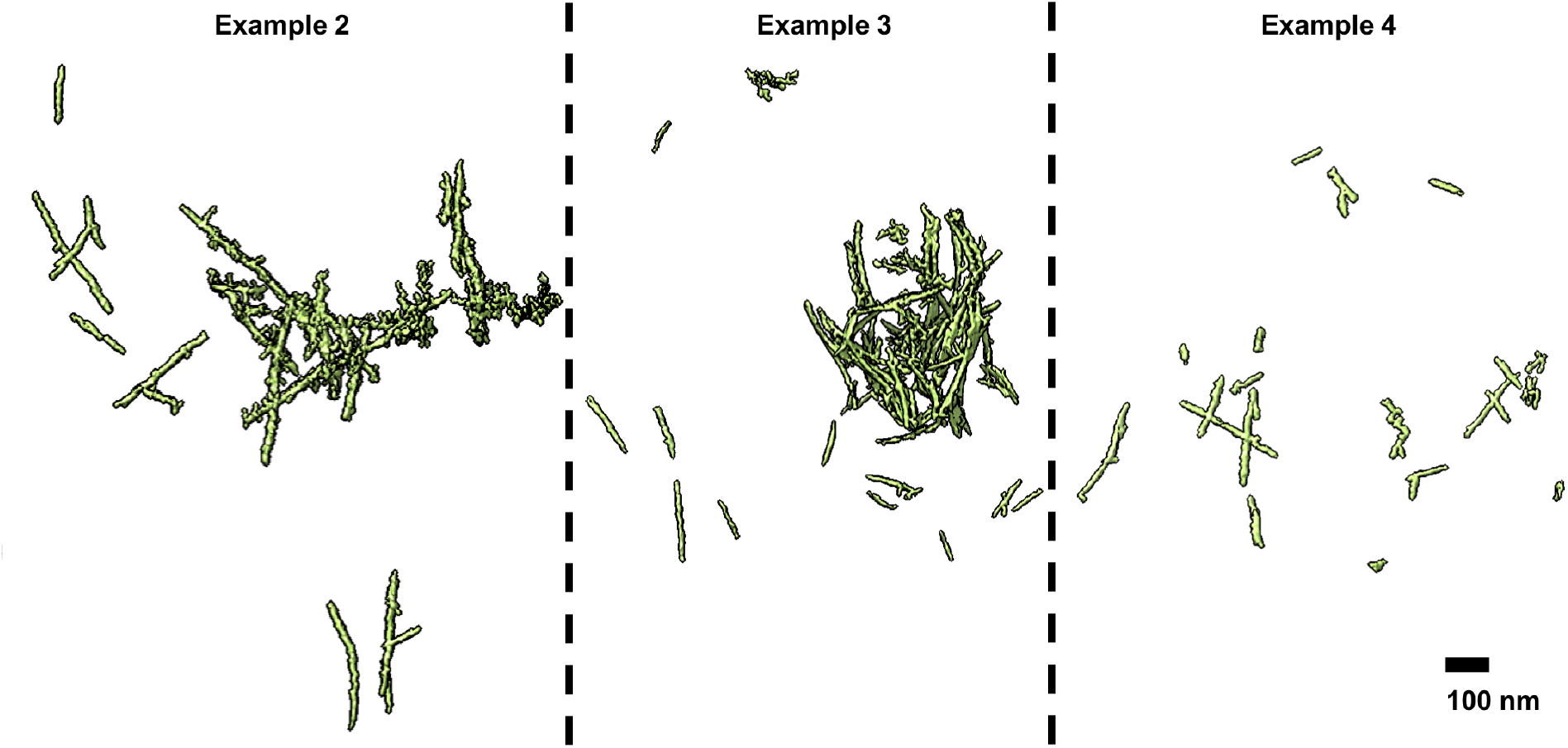
Additional examples of Q51 filamentous aggregates.

**Supplementary Figure S4.**
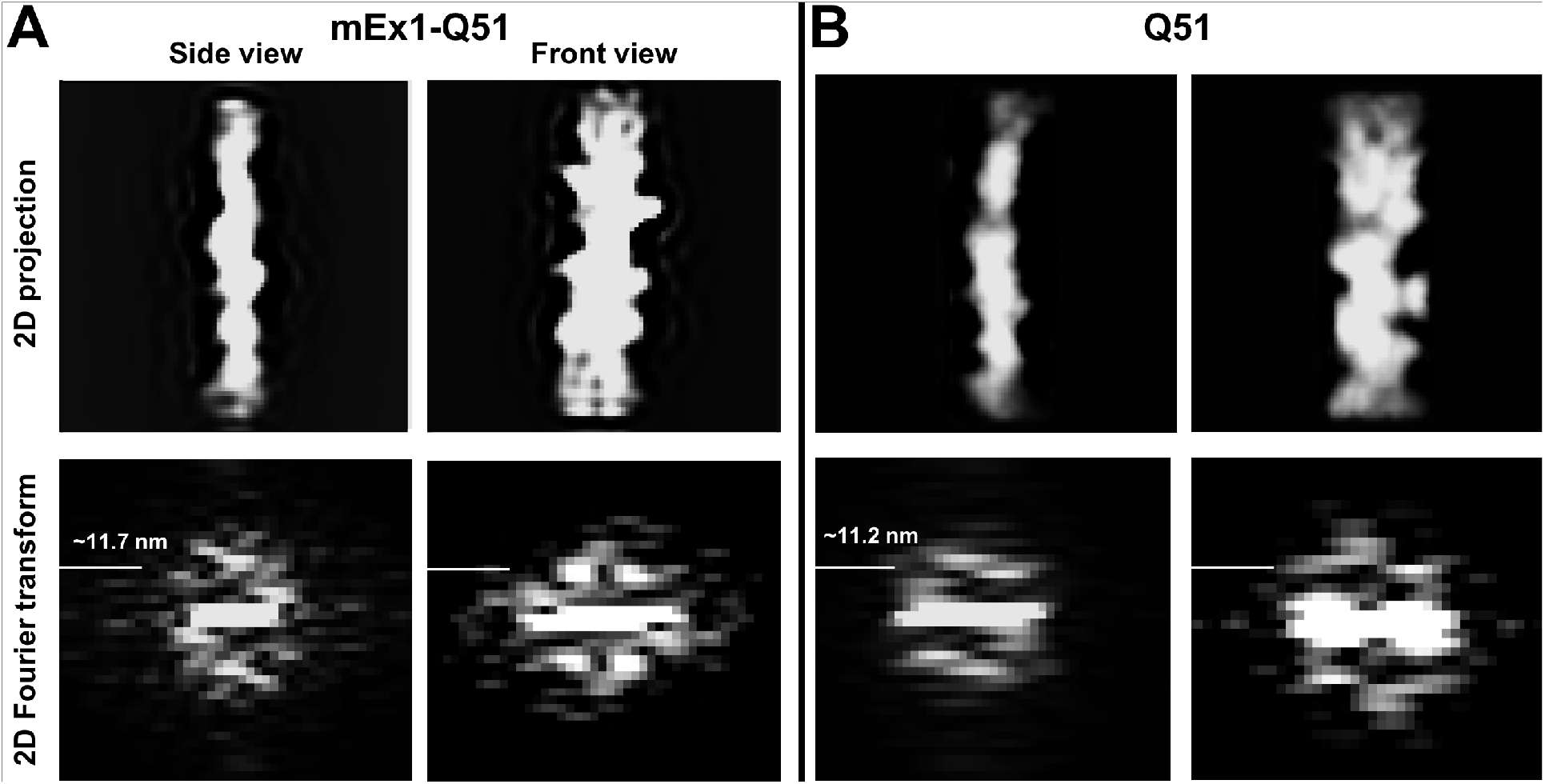
Power spectrum of orthogonal side (left) and face-on (right) projections from the subtomogram average of **(A)** mEx1-Q51 (Figure 2C) and **(B)** Q51 (Figure 4C) filaments.

